# Lessening of porcine epidemic diarrhoea virus susceptibility in piglets after editing of the CMP-N-glycolylneuraminic acid hydroxylase gene with CRISPR/Cas9 to nullify N-glycolylneuraminic acid expression

**DOI:** 10.1101/488510

**Authors:** Ching-Fu Tu, Chin-kai Chuang, Kai-Hsuan Hsiao, Chien-Hong Chen, Chie-Min Chen, Su-Hei Peng, Yu-Hsiu Su, Ming-Tang Chiou, Chon-Ho Yen, Shau-Wen Hung, Tien.-Shuh. Yang, Chuan-Mu Chen

**Affiliations:** Division of Animal Technology, Animal Technology Laboratories, Agricultural Technology Research Institute, No.1, Ln. 51, Dahu Rd., Xiangshan Dist., Hsinchu City 30093, Taiwan, R.O.C.; Chao Kun Biotech Ltd. No.22, Lane 156, Tai-Yuan Rd., Taipei 103, Taiwan, R.O.C.; Department of Veterinary Medicine, College of Veterinary Medicine, National of Science and Technology, Pingtung, Taiwan, ROC.; Division of Animal Resources, Animal Technology Laboratories, Agricultural Technology Research Institute, No.1, Ln. 51, Dahu Rd., Xiangshan Dist., Hsinchu City 30093, Taiwan, R.O.C.; Department of Life Sciences, National Chung Hsing University, No.145, Xingda Rd., South Dist., Taichung City 402, Taiwan, R.O.C.; Reproductive Medicine Center, Lee Women’s Hospital, Taichung City, Taiwan, R.O.C.; Department of Biotechnology and Animal Science, National Ilan University, No.1, Sec. 1, Shennong Rd., Yilan City, Yilan County 26047 O.C., Taiwan, R.O.C.; The iEGG and Animal Biotechnology Center, National Chung Hsinh University, Taichung 402, Taiwan, R.O.C.

## Abstract

Porcine epidemic diarrhoea virus (PEDV) devastates the health of piglets but may not infect piglets whose CMP-N-glycolylneuraminic acid hydroxylase (CMAH) gene mutated (knockouts, KO) by using CRISPR/Cas9 gene editing techniques. This hypothesis was tested by using KO piglets challenged with PEDV. Two single-guide RNAs targeting the CMAH gene and Cas9 mRNA were microinjected into the cytoplasm of newly fertilized eggs; 4 live founders generated and proven to be biallelic KO, lacking detectable N-glycolylneuraminic acid (NGNA). The founders were bred, and homozygous offspring were obtained. Two-day-old (in exps. I and III) and 3-day-old (in exp. II) KO and wild-type (WT) piglets were inoculated with TCID_50_ 1×10^3^ PEDV and then fed 20 mL of infant formula (in exps. I and II) or sow’s colostrum (in exp. III) every 4 hours. In exp. III, the colostrum was offered 6 times and was then replaced with Ringer/5% glucose solution. At 72 hours post-PEDV inoculation (hpi), the animals were euthanized for necropsy, and their intestines were sampled. In all 3 experiments, the piglets showed apparent outward clinical manifestations suggesting that infection occurred despite the CMAH KO. In exp. I, all 6 WT piglets and only 1 of 6 KO piglets died at 72 hpi. Histopathology and immunofluorescence staining showed that the villus epithelial cells of WT piglets were severely exfoliated, but only moderate exfoliation and enterocyte vacuolization was observed in KO piglets. In exp. II, delayed clinical symptoms appeared, yet the immunofluorescence staining/histopathologic inspection (I/H) scores of the two groups differed little. In exp. III, the animals exhibited clinical and pathological signs after inoculation similar to those in exp. II. These results suggest that porcine CMAH KO with nullified NGNA expression are not immune to PEDV but that this KO may lessen the severity of the infection and delay its occurrence.

**Author summary:** The infection of villus epithelial cells by PEDV has been suggested to occur via putative sialic acid and aminopeptidase N (APN) receptors. Thus, CMP-N-glycolylneuraminic acid hydroxylase (CMAH) gene-mutated pigs that lack N-glycolylneuraminic acid (NGNA) receptors should exhibit resistance to PEDV infection even when APN, which is also responsible for peptide digestion and amino acid absorption and should not be tackled and remains intact. This hypothesis was tested in the present study by generating animals of this type; however, after PEDV challenge, they still showed clinical manifestations of infection. Although the hypothesis could not be verified by the results of the study, some of the immunological and histopathological evidence obtained suggested that this genetic alteration may lessen the severity of infection and delay its occurrence. The results also suggested that binding to NGNA is not a sufficient and necessary condition for PEDV infection of enterocytes. The null expression of CMAH by gene editing induced insignificant resistance to PEDV infection in neonatal piglets.

## Introduction

Porcine epidemic diarrhoea (PED) was first recognized as an enteric disease in 1971 by the British veterinarian Oldham [*cf.* 1]; subsequently, the PED virus (PEDV) was isolated by Pensaert and de Bouck [2] at Ghent University in Belgium. Since then, PEDV-associated diarrhoea has been widely detected in Europe. In Asia, it was reported in 1982 [3], and it has subsequently greatly impacted the Asian pork industry. In China during 2010 and 2011, over one million nursing piglets were lost due to PEDV-associated diarrhoea [4]; in 2013, PEDV emerged in Korea and the USA [5–7] as well as in Taiwan [8], causing great economic losses and continuing to spread as an epidemic.

PEDV and transmissible gastroenteritis virus (TGEV) are members of the *Coronaviridae* family and the alpha coronavirus group. The PEDV genome consists of a positive single-stranded RNA approximately 28 kb in length that contains 7 open reading frames (ORF), including ORF1a, ORF1b, and ORF2-6 [9]. The viral particles are coated with S-protein, a type I membrane protein; it forms spikes on the viral surface that are used to infect host cells and also bears highly antigenic domains and could theoretically be used to develop a high-titre neutralizing PEDV vaccine [6, 10]. However, Sun et al. [11] found that the sequence of this region is highly variable, a characteristic that is likely to reduce the efficiency of conventional commercial vaccines. Furthermore, the S-protein is a glycoprotein that undergoes complicated post-translational modifications that result in antigen diversity and create obstacles to the development of a PEDV vaccine [10].

The pathway of PEDV infection occurs mainly through the S-protein. PEDV first contacts sialic acids (neuraminic acid, NA) in host intestine [12] and then infects the villi by binding to aminopeptidase N (APN) on epithelial cells [13, 14]. These findings suggest that NA is the first glycoprotein receptor and that APN is the second receptor for PEDV during infection of the host intestine [15]. A similar process occurs during infection by transmissible gastroenteritis virus (TGEV) [12]; on the other hand, porcine respiratory coronavirus (PRCV) loses its ability to infect the host intestine due to mutation and deletion of the S-protein genomic region occur [16]. Since viral genomic sequences of S-protein are generally variable and unstable, but in mammals, e.g., pigs, the codon sequences of their receptor are more stable and allow to be manipulated specifically by GE. As mentioned above, PEDV infects the host via NA and APN, and NA has been shown to play an important role in host immune function and infection by pathogens [17, 18]. Human cells are able to synthesize N-acetylNA (NANA) but not N-glycolylNA (NGNA) [19] because the human CMP-N-glycolylneuraminic acid hydroxylase (CMAH) gene has mutated during 2.5-3 million years of evolution [17]. We suggest that, analogous to the way in which human evolution has eliminated the NGNA receptor for PEDV, the CMAH gene of domestic pigs might be artificially mutated by gene editing technology to produce resistance to PEDV infection. The APN gene is not proposed as a target because it is essential for dipeptide digestion and amino acid absorption.

Currently available technologies for gene editing (GE) include the use of ZFN (zinc finger nuclease) [20], TALEN (transcription activator-like effector nuclease) [21], and CRISPR (clustered regularly interspaced short palindromic repeat)/Cas9 (CRISPR-associated (Cas) endoribonuclease 9) [22]. Due to the availability of convenient techniques for constructing and editing vectors and the fact that Cas9 is a universal enzyme that can be constructed separately to guide/target vectors, use of the CRISPR/Cas9 system for GE is currently more popular than use of the ZFN and TALEN systems. Furthermore, GE can be simultaneously conducted on multiple sites or genes with the same Cas9 to achieve different targeting purposes or reduce the risks of off-targeting [23–25]. We have established TALEN [26] and CRISPR/Cas9 [27, 28] systems for direct microinjection of GE vectors to generate GGTA1 mutant pigs. In this study, direct microinjection of two single-guide RNA and Cas9 mRNA vectors into the cytoplasm of pronuclear porcine embryos was used to generate CMAH mutant pigs with null expression of NGNA, and the possibility of obtaining mutant piglets that are resistant to infection by PEDV was examined.

## Results

### Generation of CMAH mutant pigs

A total of 67 embryos were microinjected with the CRISPR RNA, including two sgRNA which are directed against two sites on CMAH within exon2 and intron 2 (Fig 1A), and Cas9 mRNA and transferred to 3 foster dams. Five live piglets and 1 stillborn piglet were delivered by one pregnant sow (Table 1). PCR analysis revealed that 1 male (L667-02) and 3 females (L667-10, -11, and -12) (Fig 1C) carried 161-bp deletion mutations (Fig 1B). Further analysis by PCR-directive sequencing (PDS) and subcloning of PCR products in T-A cloning vectors and sequencing (PTS) showed that the 4 live piglets and the stillborn piglet were biallelic CMAH mutants; of these, L667-02 was biallelic 161-bp deleted (D/D type) (Fig 2A), L667-10, -11 and -12 were mosaic with D/D type and 2 sites mutated (D/D and D/M types) (Fig 2A-C), and the stillborn animal (L667-D) had a single base mutation, a 5-bp insertion at site I and a 5-bp deletion at site II (M/M type) (Fig 2B and C). The mutational status of their offspring (Table 2) was confirmed by PCR, PDS and PTS (S1-S3 Figs). The animals used for PEDV challenge were obtained by breeding the three founders with the founder boar. All piglets were rapidly screened by PCR, and D/D piglets were preferentially used in the experiments. In exps. II and III, the D/D piglets were supplemented with 1 and 3 D/M type piglets, respectively (Table 3) that were confirmed by PTS to have 1-bp insertions or 14- or 2-bp deletions at site I on the mutated chromosome (S1-S3 Figs). The null expression of the CMAH gene was analysed based on the detection of NGNA/NANA by HPLC; the results showed that all founders (Fig 3) and their offspring (Fig S4) lacked NGNA expression. These results show that all founders and their offspring are biallelic mutants that fail to express CMAH and produce no NGNA in their tissues.

**Fig 1.**
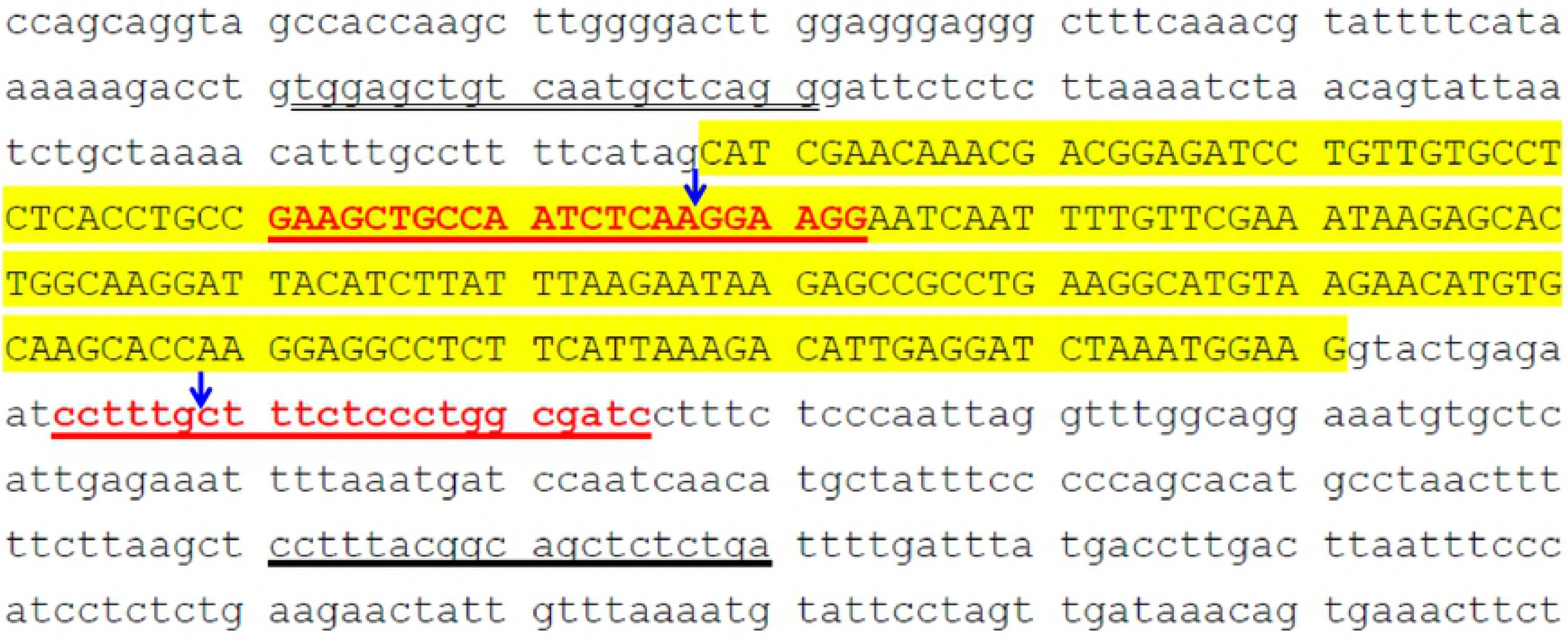
Generation of CMAH Gene-Edited Pigs. A. The porcine CMAH gene editing sites were designated on exon 2 (sense strand, red capital letters underlined in red) and intron 2 (antisense strand, red letters underlined in red). The sequences underlined in black are PCR primers. The sequences shown in large capital letters with yellow shading are exon 2. The blue arrows indicate the gene editing sites. B. CMAH KO piglets were analysed and screened by PCR. C. Four lines of CMAH gene-edited piglets (1 male and 3 females) were obtained.

**Fig 2.**
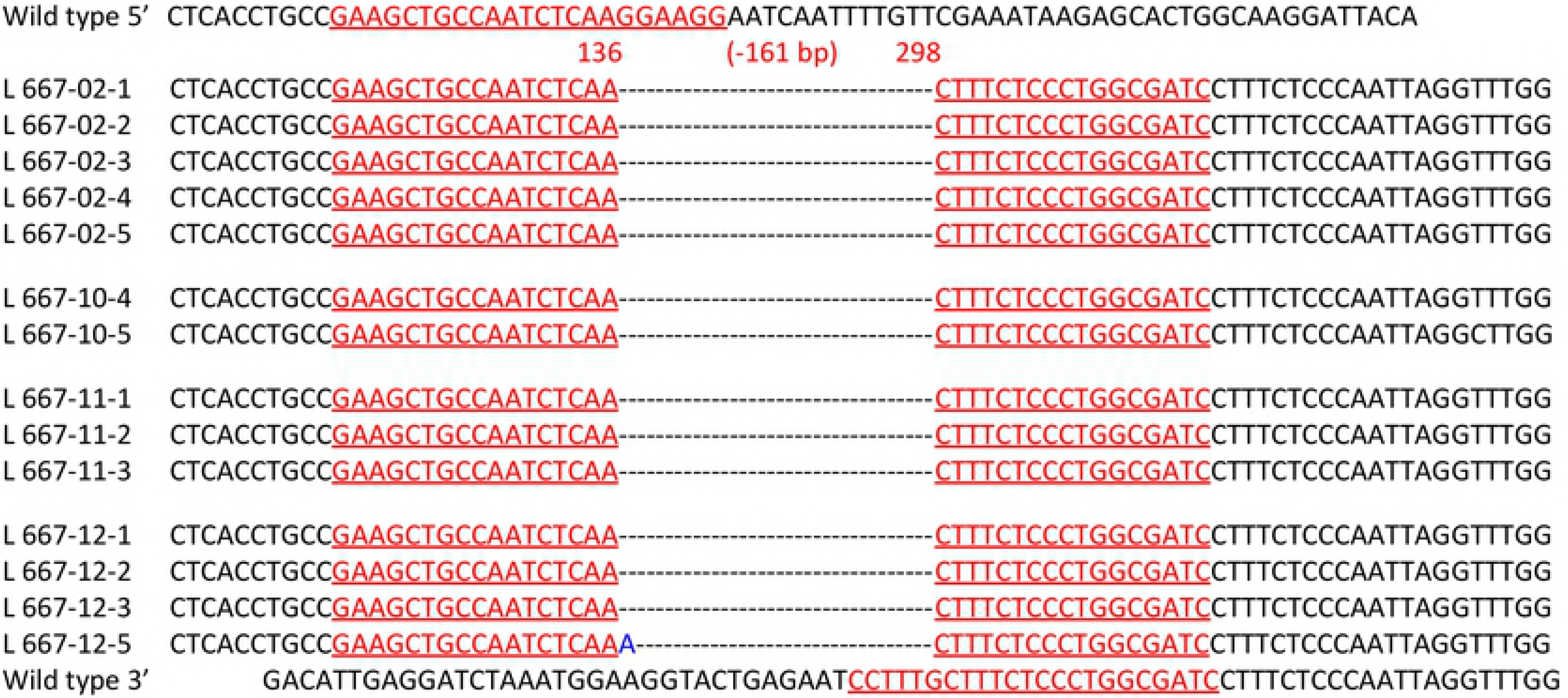
Genotyping by TA-Cloning and Sequencing of the Porcine CMAH Gene Edited by CRISPR/Cas9 Vectors Directed Against Two Sites. A. The genotype shown displays two simultaneously mutated sites and a deleted 161-bp DNA fragment; the blue A represents an extra inserted base that appeared in L667-12. B. The indel occurred at site I of exon II of the CMAH gene. C. Details of the mutation at site II of intron 2 of the CMAH gene. The blue letters represent inserted bases, and the dashed line indicates deleted bases.

**Fig 3.**
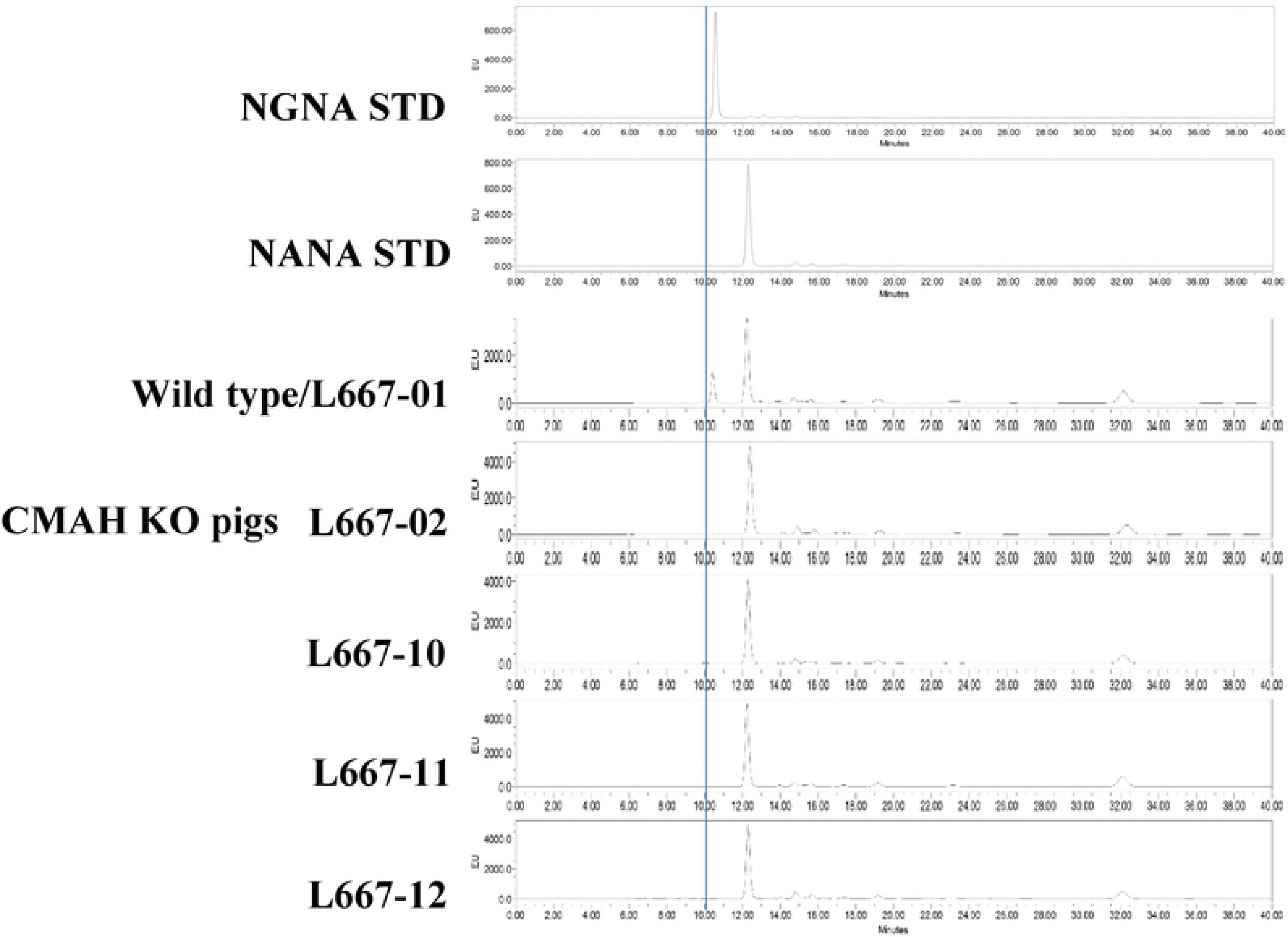
Expression of NGNA/NANA in the Tissues of CRISPR/Cas9 CMAH Mutant Founders. L667-02, -10, -11 and -12 and their wild-type littermate (L667-01) were analysed by HPLC. NGNA STD and NANA STD are standard samples of NGNA and NANA, respectively. The line indicates a retention time of 10 min.

**Table 1.** Generation of CMAH Knockout (KO) Pigs by Direct Microinjection of sgRNA/Cas9 mRNA into the Cytoplasm of Pronuclear Newly Fertilized Porcine Eggs.

| Micromanipulation |  | Surrogate dam |  | No. of piglets |  |  |
| --- | --- | --- | --- | --- | --- | --- |
| Exp. | No. Embryos | E.T. | Pregnant (%) | Born<sup>a</sup> | KO (%) | BKO<sup>b</sup> (%) |
| 1 | 31 | 1 | 1 | 5/1 | 5 (83.3) | 5 (83.3) |
| 2 | 21 | 1 | 0 | 0/0 | 0 (0) | 0 (0) |
|  | 18 | 1 | 0 | 0/0 | 0 (0) | 0 (0) |
| Total | 67 | 3 | 1 (33.3) | 5/1 | 5 (83.3) | 5 (83.3) |

<sup>a</sup>. alive/dead
<sup>b</sup>. biallelic knockout

**Table 2.**
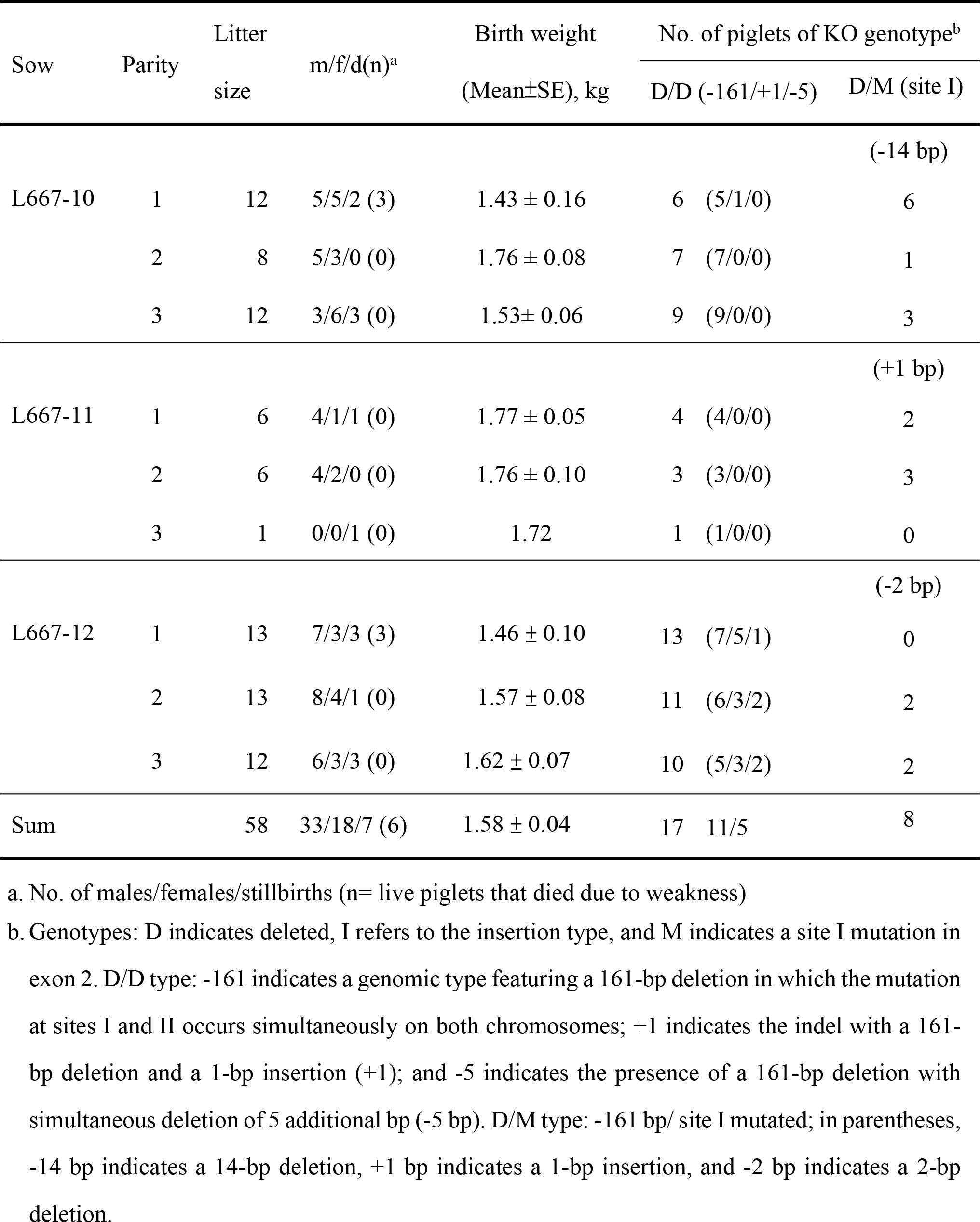
Germline Transmission and Genotypes of F1 CMAH KO Piglets.

**Table 3.** Genotypes of the F1 CMAH KO Piglets Used for PEDV Challenge.

| Exp. | Sow/ L667- |  |  | Genotype |  | Site I of<br>Mutant allele |
| --- | --- | --- | --- | --- | --- | --- |
|  | Parity | ID. | Litter size | D/D | D/M |  |
| 1 | 1 | 10 | 12 | 1 | 0 | - |
|  |  | 11 | 6 | 2 | 0 | - |
|  |  | 12 | 13 | 3 | 0 | - |
| Control |  | 12 |  | 1 | 0 | - |
| 2 | 2 | 10 | 8 | 3 | 0 | - |
|  |  | 11 | 6 | 3 | 1 | +1 |
|  |  | 12 | 13 | 2 | 0 | - |
| Control |  | 12 |  | 1 | 0 | - |
| 3 | 3 | 10 | 12 | 4 | 2 | -14 |
|  |  | 11 | 1 | 0 | 0 | - |
|  |  | 12 | 12 | 5 | 1 | -2 |
| Control |  | 10 |  | 1 | 0 | - |

Note: Genotypes: D indicates that the mutation occurred at two sites simultaneously and resulted in a 161-bp deletion, whereas M is site I-mutated; D/D refers to the case in which the 161-bp deletion occurred on both chromosomes. The controls are wild-type piglets.

### Clinical observation of neonatal piglets challenged with nv-PEDV

#### Exp. I

When neonatal 2-day-old piglets were challenged with nv-PEDV, both the CMAH mutant (Knockout, KO) and wild-type (WT) animals initially displayed clinical signs of vomiting and diarrhoea at 12 hours post-inoculation (hpi), and their activity also decreased (Table 4). In the WT group, the first piglet’s death occurred at 44 hpi; a second animal died at 52 hpi, a third at 68 hpi, and the remaining three animals were moribund and nearly dead at 72 hpi (Fig 4A). In the CMAH KO group, the first piglet died at 60 hpi, 3 piglets were moribund at 72 hpi, and the other remaining two piglets survived until the end of the trial (Fig 4A and Table 4). After nv-PEDV inoculation, the loss of body weight of WT piglets was 0.69±0.04 kg, significantly (p<0.01) greater than that of CMAH KO piglets (0.45±0.03 kg) (Fig 5A).

**Table 4.** Clinical Signs Displayed by Neonatal Piglets after nv-PEDV Inoculation in Exps. I and II.

| Exp. | Geno-type | No. | Hours post nv-PEDV inoculation |  |  |  |  |  |  |
| --- | --- | --- | --- | --- | --- | --- | --- | --- | --- |
|  |  |  | 4-8 | 12 | 24 | 36 | 48 | 60 | 72 |
| I | KO | 6 | 6A/ | 6B/ | 6B/ | 1A5B/ | 6B/ | 4B1C1D/ | 2B3C1D/ |
|  |  |  | 6n | 2d4dv | 3n3d | 4n2d | 3n3d | 1n4d | 5d |
|  | WT | 6 | 6A/ | 6B/ | 2B4C/ | 4B1C1D/ | 4B1C1D/ | 1B3C2D/ | 3C3D/ |
|  |  |  | 6n | 6dv | 2n4d | 6d | 2n2d1dv | 4d | 3d |
| II | KO | 9 | 9A/ | 9A/ | 9B/ | 9B/ | 6B2C1D/ | 1B1C7D | 9D/ |
|  |  |  | 9n | 2n1v2d4dv | 2n6d1dv | 9d | 8d | 2d | 0 |
|  | WT | 9 | 9A/ | 9A/ | 9B/ | 9B/ | 5B2C2D/ | 2B7D/ | 1B1C7D/ |
|  |  |  | 9n | 4n3v1d1dv | 1n8d | 1n8d | 7d | 2d | 2d |

\* No. of piglets with viability and clinical signs: viability - A is normal, B indicates decreased activity, C is moribund, and D indicates dead; clinical signs - n indicates no clinical signs, d is diarrhoea, and v is vomiting.

**Fig 4.**
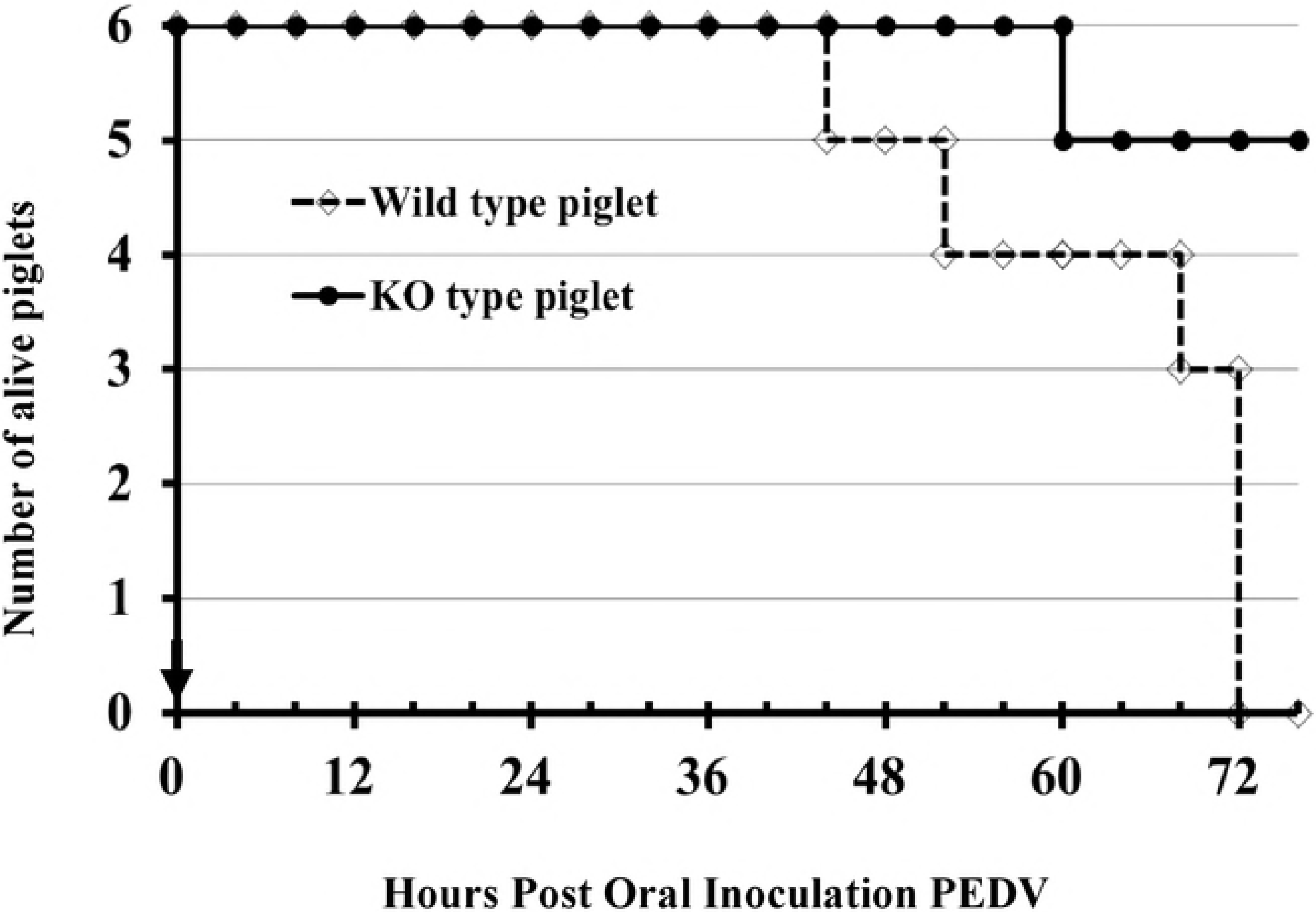
Survival of Neonatal Piglets After Oral Inoculation with nv-PEDV. A, 2-day-old piglets; B, 3-day-old neonatal piglets inoculated with PEDV. Solid circles (A) or squares (B) with lines represent the CMAH KO piglets, and open diamonds (A) or squares (B) with dashed lines indicate wild-type piglets. The arrow shows the time of inoculation. In A at 72 hpi, three moribund WT piglets are classified as dead piglets.

**Fig 5.**
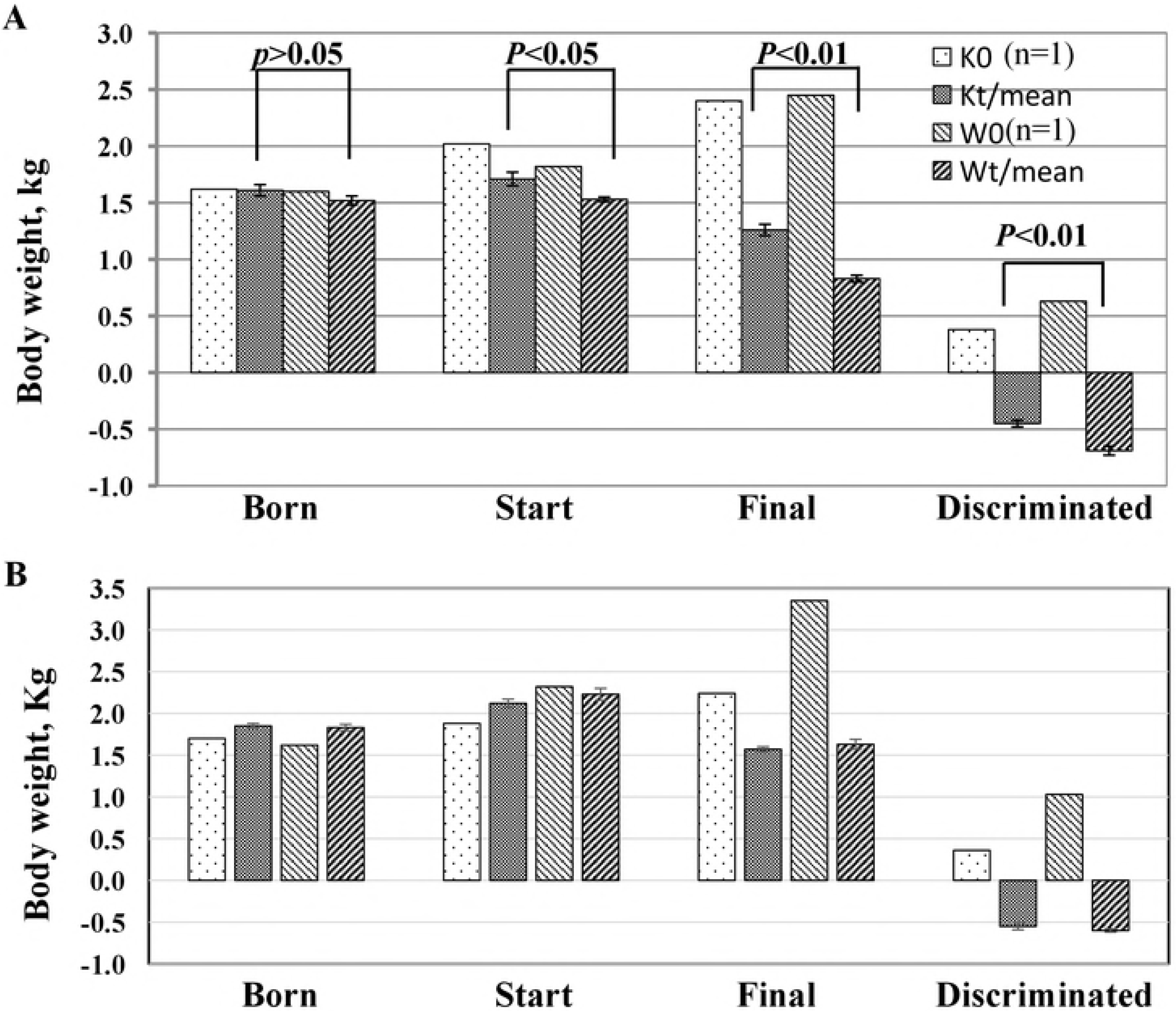
Body Weights of Neonatal Piglets Before and After Oral PEDV Inoculation. A and B show 2-day-old piglets (n=6) and 3-day-old piglets (n=9), respectively. K0 and W0 represent KO and WT animals that were not inoculated with PEDV and were reared by their dams on the farm. KO and WT are knockout treated and wild-type treated animals, respectively.

#### Exp. II

The 3-day-old piglets were examined as in exp. I. Although both CMAH KO and WT animals initially showed clinical signs of vomiting and diarrhoea at 12 hpi, 2 KO and 4 WT piglets were without clinical signs (Table 4). Furthermore, all piglets sustained their activity until 24 hpi (Table 4). In the WT group, the first death occurred at 40 hpi (Fig 4B); two piglets were lost at 48 hpi, 4 piglets died at 56 hpi, and the remaining two piglets were alive at the end of the trial. In the CMAH KO group, the first animal was lost at 44 hpi, and 3, 3, 1 and 1 piglets died at 52, 56, 64 and 68 hpi, respectively (Fig 4B). There was no significant difference in the decrease in body weight in the two groups of piglets (WT/ −0.60±0.02 kg vs. CMAH KO/-0.55 ±0.04 kg; p>0.05) (Fig 5B).

#### Exp. III

To examine the early events and the role of NGNA in nv-PEDV infection of neonatal piglets, we used 2-day-old piglets challenged with nv-PEDV. After infection, the piglets were fed sows’ milk and skim milk every 4 hours for 24 hours; this was then replaced by Ringer’s lactate solution supplemented with 5% glucose, and the piglets were sacrificed at 24, 48 and 72 hpi. The results (Table 5) show that until 12 hpi both the CMAH KO and WT piglets appeared normally active; however, with respect to clinical signs, only 3/11 CMAH KO piglets did not show diarrhoea or vomiting at 12 hpi. From 4 to 24 hpi, all piglets were fed their own dams’ whole or skim milk; the results show that all piglets displayed decreased activity and diarrhoea without a significant difference between CMAH KO and WT piglets. One moribund CMAH KO piglet was observed at 44 hpi, and one moribund WT piglet was observed at 56 hpi; all of the piglets stopped vomiting after 24 hpi. After the sow’s milk was replaced with RLG, all piglets (both CMAH KO and WT) showed sustained activity and viability at least until 56 hpi, with the exception of one CMAH KO piglet that died prior to the end of the trial (Table 5).

**Table 5.** Clinical Signs Displayed by Neonatal Piglets after nv-PEDV Inoculation in Exp. III.

| Geno-<br>type | h | No. | Hours post nv-PEDV inoculation |  |  |  |  |  |  |  |  |  |  |
| --- | --- | --- | --- | --- | --- | --- | --- | --- | --- | --- | --- | --- | --- |
|  |  |  | 4-8 | 12 | 16 | 20 | 24 | 28-40 | 44 | 48 | 56 | 64 | 72 |
| KO | 24 | 3 | 3A/<br>3n | 3A/<br>2n1v | 3B/<br>3d | 2B1C/<br>1v2d | 3B/<br>3d | - | - | - | - | - | - |
|  | 48 | 2<sup>#</sup> | 2A/<br>2n | 2A/<br>1n1v | 2B/<br>2d | 2B/<br>1d1dv | 2B/<br>2d | 2B/<br>2d | 1B1C/<br>1n1d | 2B/<br>1n1d | - | - | - |
|  | 72 | 6 | 6A/<br>6n | 6A/<br>4v1d1dv | 6B/<br>6d | 6B/<br>6d | 6B/<br>6d | 6B/<br>6d | 6B/<br>6d | 6B/<br>6d | 4B2C/<br>6d | 3B3C/<br>6d | 3B2C1D/<br>5d |
| WT | 24 | 3 | 3A/<br>3n | 3A/<br>2v1dv | 3B/<br>3d | 2B/<br>2d1dv | 3B/<br>3d | - | - | - | - | - | - |
|  | 48 | 3 | 3A/<br>3n | 3A/<br>1v1d1dv | 3B/<br>3d | 3B/<br>3d | 3B/<br>3d | 3B/<br>3d | 3B/<br>3d | 3B/<br>3d | - | - | - |
|  | 72 | 6 | 6A/<br>6n | 6A/<br>2v4dv | 6B/<br>6d | 6B/<br>6d | 6B/<br>6d | 6B/<br>6d | 6B/<br>1n5d | 6B/<br>6d | 5B1C/<br>6d | 4B2C/<br>6d | 5B1C/<br>6d |

\* No. of piglets with viability and clinical signs: viability - A is normal, B indicates decreased ability, C is moribund, and D is dead; clinical signs - n indicates no clinical signs, d is diarrhoea, and v is vomiting.
<sup>#</sup> One piglet died before experiment.

### Immuno/histopathology of neonatal piglets challenged with nv-PEDV

After 72 hpi, all of the dead and euthanized piglets were necropsied, and their intestines were sampled for pathological examination. Grossly, the small intestine appeared transparent and orange-yellow to flesh pink in colour; it was thin-walled and dilated with fluid content in the live piglets (S5A-B Fig). In exp. I, the PEDV induced histopathologic changes, including enterocyte necrosis, degeneration, and exfoliation, and collapsed lamina proprial tissues containing karyorrhectic debris, were noted in all challenged piglets. However, these lesions varied from mild to severe, and the lesions were more severe in the moribund WT piglets than in the CMAH KO piglets (Fig 6). Immunofluorescence (IF) staining with a monoclonal antibody against PEDV nuclear protein was used to detect PEDV antigens. The results showed that PEDV antigen was presented in the epithelium covering the moderately atrophic tips of villi in the small intestine of WT and CMAH KO piglets (Fig 7). However, if the epithelial cells were defoliated from the villi after PEDV infection, no positive signals would be expected (Fig 7A). We further scored the severity of lesions in the intestines of the infected animals (Fig 8) by a combination of IF staining and histopathological inspection (immuno/histopathological, I/H, score). According to the I/H scores, there appeared to be no significant difference in the severity of the intestinal lesions in WT piglets (from 3.4±0.6 to 4.4±0.3) and those in CMAH KO piglets (from 4.3±0.4 to 4.7±0.2) in exp. II (Table 6). In exp. III, even ruling out the possible effects of feeding the animals commercial baby cow milk, we also found no significant difference in the I/H scores of WT and CMAH KO piglets (Table 7). According to the I/H scores obtained at 72 hpi, which ranged from 3.8±0.4 to 3.2±0.5 in CMAH KO piglets and from 2.8±0.4 to 2.5±0.2 in WT piglets (*p*>0.05), most piglets seemed to improve compared with those at 24 and 48 hpi when offered sow’s milk and supplemental lactated Ringer’s solution containing 5% glucose (Table 7).

**Fig 6.**
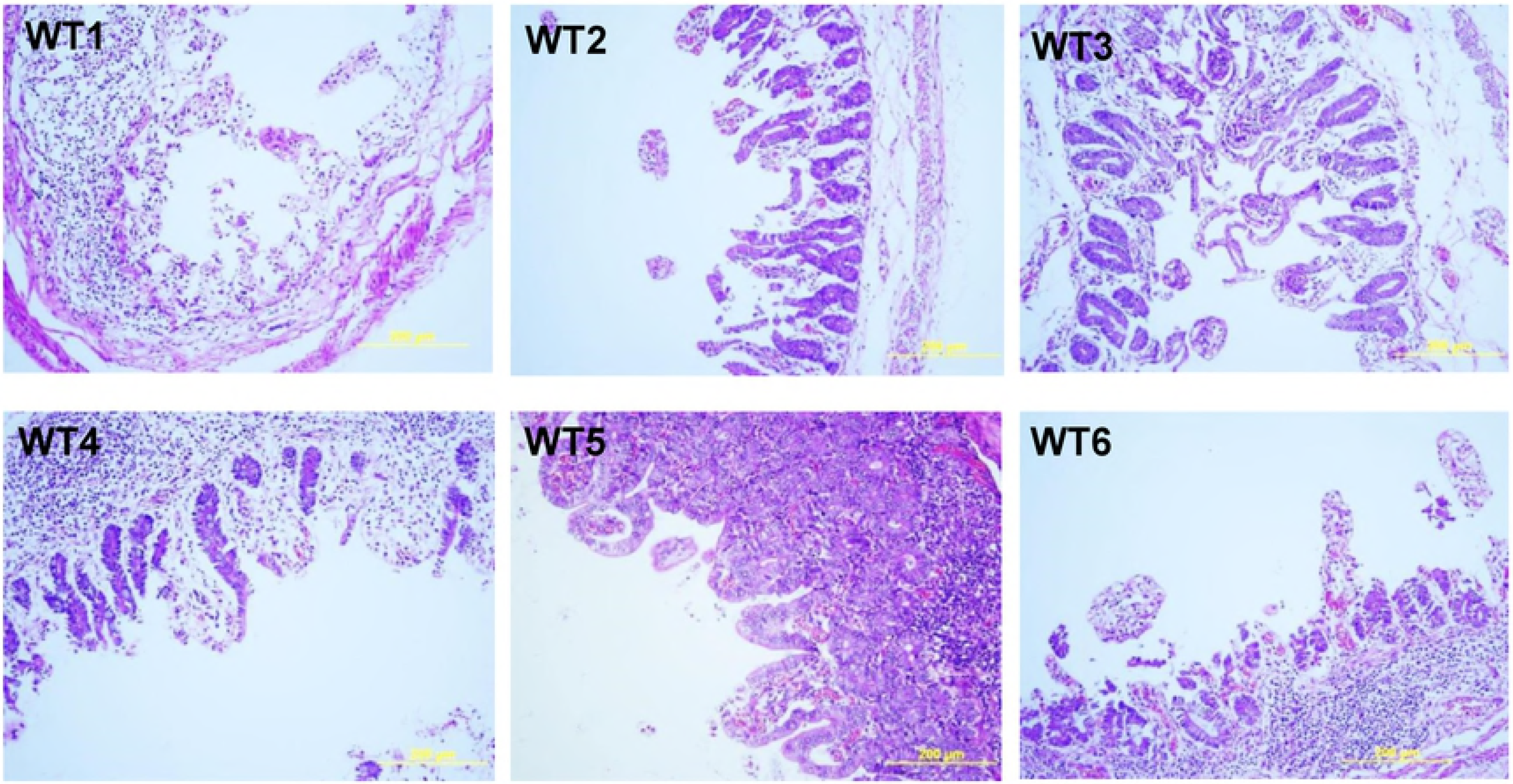
Pathological Inspection of Piglets’ Intestine at the Middle Jejunum by H/E Staining. Panels A-1 and B1 and A-2 and B2 indicate wild-type and knockout piglets, respectively, after PEDV oral inoculation. The yellow bars indicate 200 μm. The piglets from which these samples were obtained (KO2, KO4, KO5, WT2, WT5 and WT6) were moribund at 72 hpi.

**Fig 7.**
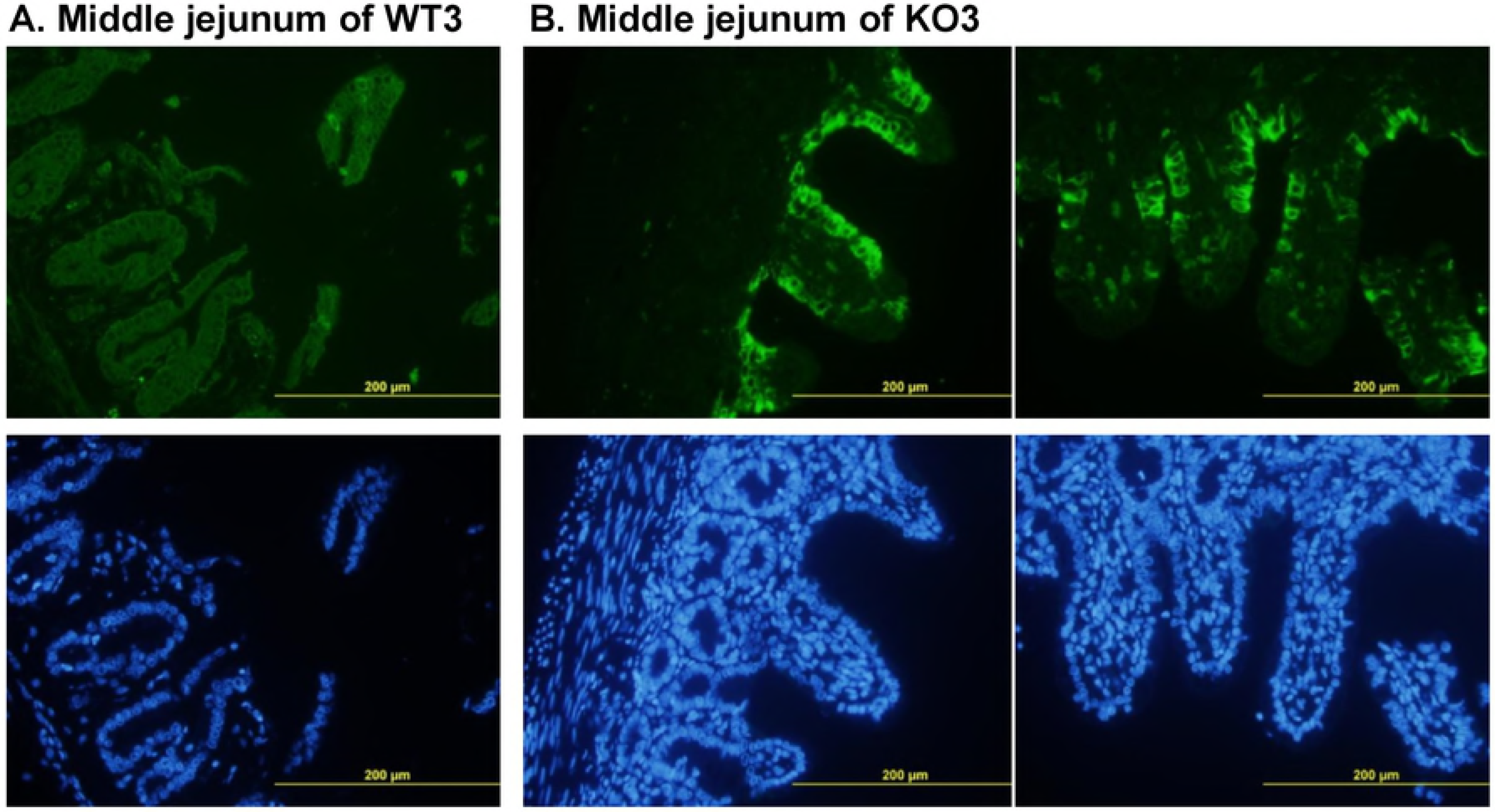
Immunofluorescence Staining with an Antibody against PEDV N Protein. WT3 (A) shows a sample from a wild-type piglet, and KO3 (B) shows a sample from a double-chromosome CMAH gene knockout piglet. The samples shown in the lower panel were stained with DAPI. The yellow bars indicate 200 μm.

**Fig 8.**
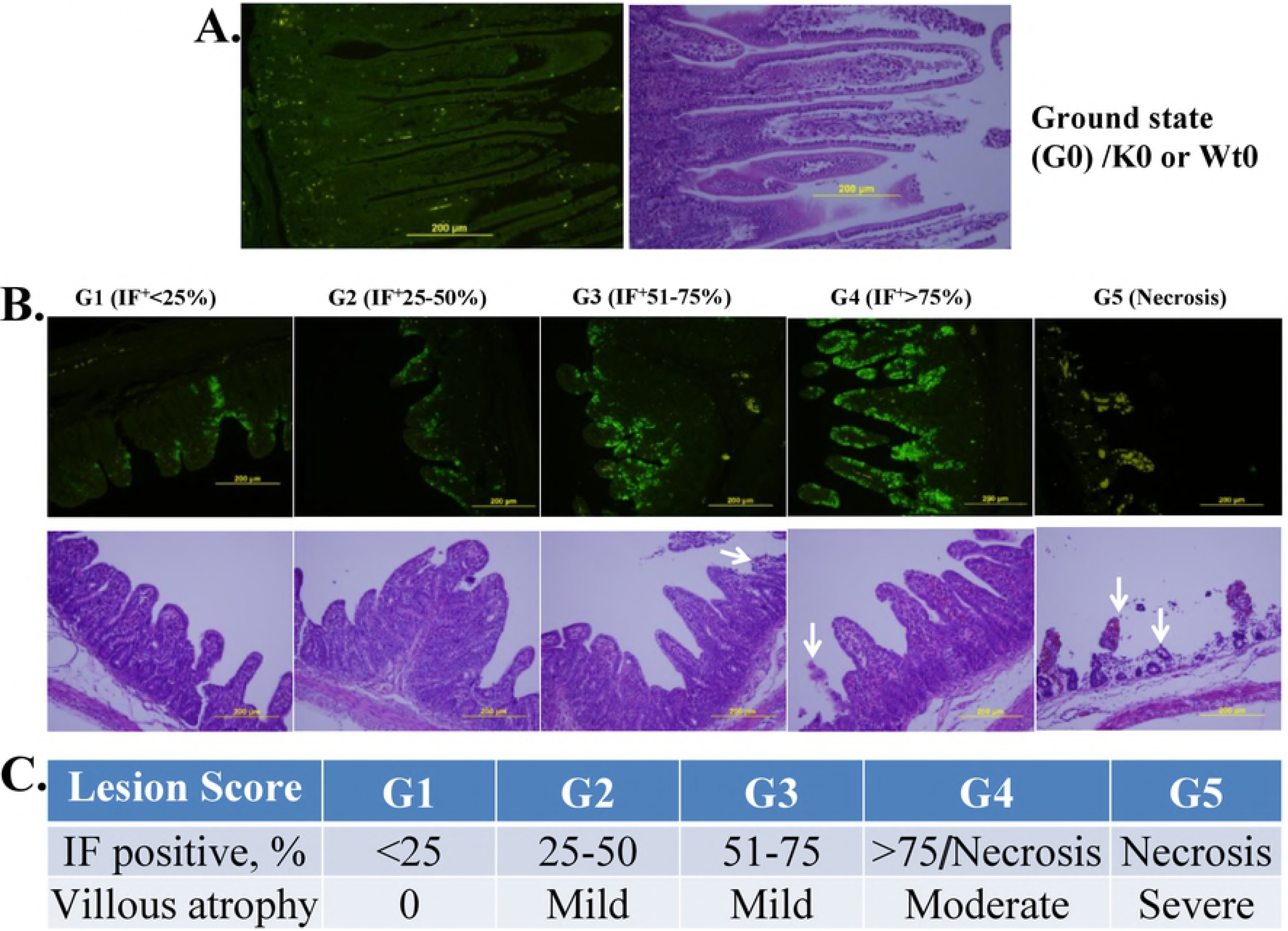
Evaluation Criteria Based on Immunofluorescence Staining and Histopathological Lesions (I/H score) of Piglets’ Intestine Samples After PEDV Challenge. A. KO0 or WT0 controls for the ground state, G0. B. IF is scored as G1 to G4 based on the relative intensity of staining, whereas G5 is based on villar atrophy or defoliation observed by H/E inspection. The corresponding scores are shown in C. The arrows indicate necrotic villi; the yellow bars represent 200 μm.

**Table 6.** Pathological Scoring of Piglet Small Intestine at 72 h after Oral Inoculation of 3-Day Old Neonates with PEDV.

| Genotype | n | Jejunum |  | Ileum |
| --- | --- | --- | --- | --- |
|  |  | Front | Middle |  |
| KO | 9 | 4.3±0.4 | 4.7±0.2 | 4.4±0.4 |
| WT | 9 | 4.4±0.3 | 3.4±0.6 | 3.6±0.6 |

KO= CMAH KO homozygotes, WT= wild-type piglets.

**Table 7.**
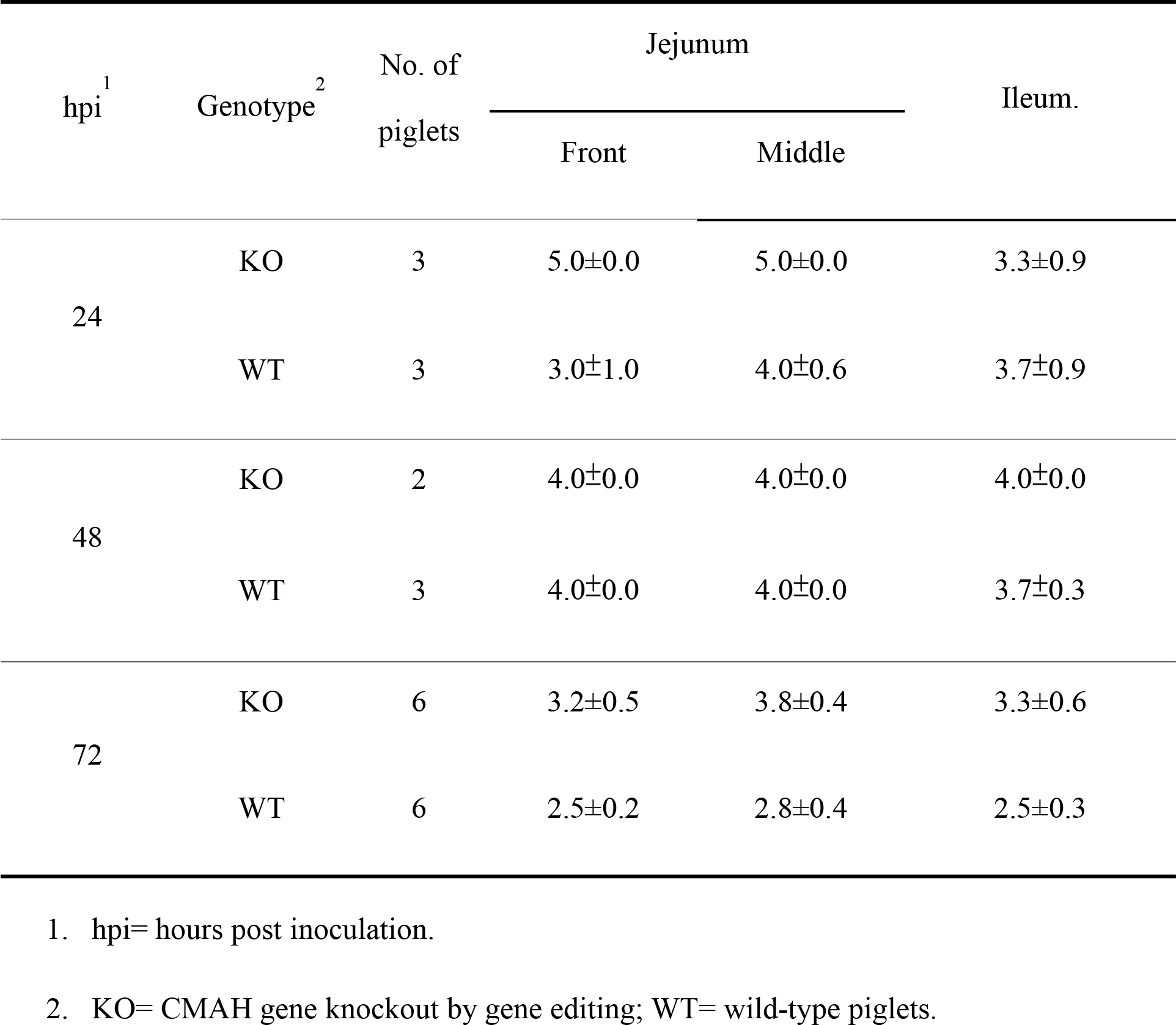
Intensity of PEDV Infection of Epithelial Cells of Villi in CMAH KO Piglets’ Intestines Revealed by Immunofluorescence Staining.

| hpi<sup>1</sup> | Genotype<sup>2</sup> | No. of piglets | Jejunum |  | Ileum. |
| --- | --- | --- | --- | --- | --- |
|  |  |  | Front | Middle |  |
| 24 | KO | 3 | 5.0±0.0 | 5.0±0.0 | 3.3±0.9 |
|  | WT | 3 | 3.0±1.0 | 4.0±0.6 | 3.7±0.9 |
| 48 | KO | 2 | 4.0±0.0 | 4.0±0.0 | 4.0±0.0 |
|  | WT | 3 | 4.0±0.0 | 4.0±0.0 | 3.7±0.3 |
| 72 | KO | 6 | 3.2±0.5 | 3.8±0.4 | 3.3±0.6 |
|  | WT | 6 | 2.5±0.2 | 2.8±0.4 | 2.5±0.3 |

1. hpi= hours post inoculation.
2. KO= CMAH gene knockout by gene editing; WT= wild-type piglets.

## Discussion

Currently, gene editing is widely used in both basic and applied studies, e.g., in studies of the disease resistance of farm animals. One convincing report showed that CD163 gene-edited pigs generated by CRISPR/Cas9 exhibited physiological normality and showed little vulnerability to porcine reproductive and respiratory syndrome virus (PRRSV) infection either in vitro [29] or in vivo [30–32]. However, other attempts, including CD169 KO and CD163 KO, failed to produce evident resistance to PRRSV [33] or African swine fever [34], respectively. These failures may have occurred because the mechanism of viral infection involves other receptors or because it does not involve receptors [35].

We have established TALEN [26] and CRISPR/Cas9 [27, 28] gene editing techniques in which the editing plasmid vector is directly microinjected into the pronucleus of newly fertilized eggs and have used these techniques to generate GGTA1 mutant pigs for the study of xenotransplantation. In this study, two sgRNAs directed against exon II and intron 2 of the CMAH gene, together with Cas9 mRNA, were microinjected into the cytoplasm near the pronucleus; the microinjected eggs yielded 4 live pigs and one stillborn pig carrying a null function of the CMAH gene. Since CRISPR/Cas9 vectors make construction of the knockout vector very convenient and achieve very powerful mutation results, only 67 embryos were used for direct cytoplasmic microinjection, and 5 mutants were successfully generated. The efficiency was 7.5% based on the number of manipulated embryos and 83.3% based on the number of delivered piglets, all of which were biallelic mutants (Table 1). However, because the CRISPR/Cas9 vector might easily generate off-targeting mutants [23–25], we used two editing sites, simultaneously deleting a short DNA fragment and facilitating mutant screening by PCR. The results revealed that all the live founders carrying 161 bp deletion, especially L667-02, the sole male, was the biallelic D/D type. Burkard et al. [29] used two sgRNA targeting sites to precisely delete exon 7 of CD163 to nullify domain 5, which is required for binding PRRSV, without affecting the animals’ normal immunology or physiology. According to animal breeding practices, it is easier to confirm the founder by PCR without a requirement for DNA sequencing; therefore, the D/D type will be used in future studies. The hypothesis that the absence of NGNA expression in CMAH KO piglets disables PEDV infection was partially proven in this study. In exp. I, 2-day-old old piglets were orally inoculated with the local outbreak strain nv-PEDV [36]. Although the final (72 hpi) survival rate differed little in the WT and KO animals, based on the histopathologic examination and considering the 3 deadly moribund WT piglets (Table 4, Fig 4A), the CMAH KO piglets showed greater resistance to nv-PEDV infection than the WT animals. This assumption is supported by the high degree of histopathologic severity found in the WT piglets (WT2, WT5 and WT6), which clearly differed from that observed in the CMAH KO piglets (KO2, KO4 and KO5) (Fig 6). However, when 3-day-old piglets were used, no differences between CMAH KO and WT piglets were observed (Table 4, Fig 4B). It is doubtful that the NGNA present in cow’s milk-based formula would enable the virus to infect the CMAH KO piglets. In exp. III, colostrum from the KO or WT sows was given to avoid any possible NGNA inference, yet the final susceptibilities of the two genotypes were similar. However, at least 3 of the 11 CMAH KO piglets showed normal activity and no clinical signs (no vomiting or diarrhoea) at 12 hpi, whereas the control piglets displayed vomiting and/or diarrhoea (Table 5). Lessened severity was therefore observed.

Considering that transmissible gastroenteritis virus (TGEV) and other coronaviruses use sialic acid (neuraminic acid, NA) as their first receptor [12, 15], PEDV might act in a similar manner. The major components of porcine mucin in the small intestinal submucosa are two types of NA, N-acetylneuraminic acid (NANA) and N-glycolylneuraminic acid (NGNA) (our unpublished data). Using a glycan screening array, Liu et al. [37] showed that Neu5Ac (or NANA) has the highest binding affinity for PEDV S1-NTD-CTD; however, they also found that porcine mucin or bovine mucin could inhibit or block in vitro PEDV and TGEV infection of PK-15 or Huh-7 cells transfected with porcine APN. The present results show that CMAH KO piglets exhibited delayed infection and minor symptoms after oral PEDV inoculation, suggesting that in CMAH KO piglets that are normally nursed, PEDV may be unable to bind efficiently to the APN on the villi of epithelial cells and pass through the intestinal lumen.

Fig 4A indicates that in exp. I six WT piglets (including 3 extremely moribund piglets) and one CMAH KO piglet died within 72 hpi. Mortality and weight loss (Fig 5A) showed significant differences (p<0.01), although the initial body weights of the WT and KO piglets also differed (p<0.05); however, the clinical outcome showed no relation to body weight. In general, the small intestines of the four KO piglets appeared normal (S5B Fig). H/E staining of the wild-type piglets showed that villus epithelial cells in the proximal and middle portions of the jejunum and ileum were severely defoliated except in the case of WT5 (Fig 6A-1 and B-1). In contrast, in the KO piglets, only KO6 (the dead piglet) showed epithelial cells severely defoliated from villi (Fig 6A-2 and B-2). However, these differences between CMAH KO and WT piglets were not observed after PEDV inoculation in exps. II and III according to the I/H scores obtained at 24, 48 and 72 hpi (Table 7). Actually, in exp. III at 24 hpi, we found that the I/H scores of CMAH KO (5.0 ±0. 0) and WT (3.0 ± 1.0) were not significantly different (p = 0.12).

It is known that PEDV causes severe enteric disease in suckling piglets [38, 39] and less severe disease in older weaned pigs [40]. Our results suggest that the differentiation might occur as early as in the neonatal period; clinical diarrhoea and/or vomiting and decreased activity were observed in all 2-day-old piglets but improved in 3-day-old piglets (Table 4). When caesarean-delivered and colostrum-deprived (CDCD) animals were used for oral inoculation of PEDV, the 1-day-old piglets showed clinical signs at 12 hpi [41]; this was also observed in our study using naturally delivered piglets. Furthermore, in PEDV inoculation studies, 5-day-old CDCD piglets were more sensitive than 21-day-old weaned piglets [42]. Similarly, naturally delivered 9-day-old suckling piglets showed a weaker innate immune response to PEDV than weaned pigs [43]. This study used 2- or 3-day-old piglets that were naturally delivered and nursed with colostrum by CMAH KO or WT sows prior to PEDV oral inoculation in an attempt to realize the protective effects of nursing in animals in which the biallelic CMAH genes were mutated. In exp. III, the clinical symptoms of 2-day-old piglets that were PEDV inoculated and hand fed whole or skim sow’s milk for an additional 24 h (Table 5) were similar to those of the 3-day-old piglets in exp. II (Table 4). Furthermore, when lactated Ringer’s solution supplemented with 5% glucose was offered from 24 to 72 hpi, the epithelial cells of the villi showed less damage and/or showed increased recovery of epithelial cells from the crypts according to the I/H scores (Table 7), which ranged from 4.0 ± 0.0 to 2.5 ± 0.2 in WT piglets and from 5.0 ± 0.0 to 3.2 ± 0.5 in the KO group. This benefit of oral rehydration therapy in acute viral diarrhoea could be attributed to glucose-facilitated sodium absorption [44]. Currently, the model may be improved by inoculating the piglets and allowed them to be continually nursed by dams of the same genotype to avoid NGNA interference.

In addition to their disease resistance, CMAH and GGTA1 KO animals are likely to display reduced hyperacute rejection of xenografts [45]. Our unpublished data also revealed that the acellular extracellular matrix derived from the intestine of CMAH KO pigs caused significantly less inflammation than that obtained from WT pigs after intramuscular implantation into CMAH/GGAT1 double KO pigs. Furthermore, NGNA present in red meat has been suggested to be a risk factor for human colorectal cancer and atherosclerosis in persons who habitually consume red meat [46]. Therefore, CMAH mutant pigs generated by GE can be viewed as pigs that offer a source of healthy red meat and of material that is suitable for use in biomedical devices.

In conclusion, the CMAH mutant pigs generated by gene editing could be a new breed with less susceptibility to PEDV, a source animal for medical materials and xenografts, and a source of healthy red meat.

## Materials and methods

### Ethics statements

All animals were managed and treated with permission from the Agricultural Technology Research Institute (IACUC104004). The use of the animals and the PEDV challenge protocol were approved by the IACUC committee of Agricultural Technology Research Institute (IACUC105063C1) and of National Pingtung University of Science and Technology (NPUST) (NPUST-105-060) under the regulation of the Animal Protection Act 1998.

### Animals and animal care

Landrace mature gilts at least 120 to 150 kg in weight or sows and their neonatal piglets were used in this study. All animals were reared in a station free from specific pathogens (atrophic rhinitis, *Mycoplasma hyopneumoniae*, pseudorabies, *Actinobacillus pleuropneumoniae*, swine dysentery, scabies, classical swine fever, foot and mouth disease and porcine reproductive and respiratory syndrome). The gilts or sows were housed indoors on concrete floors, and the accommodation was artificially lit (450-600 lux for 9 hours a day) and exposed to window sun light. The animals were fed a restricted (4% body weight) commercial diet formulated to meet the requirements recommended by the National Research Council [47] and had *ad libitum* access to water.

### Treatment of donors and recipients

The donors were synchronized and induced to super-ovulate by being fed a ration supplemented with Regumate^®^ (containing 0.4% Altrenogest; Intervet, MSD, France) for 15 days to synchronize their oestrus cycles and then being intramuscularly injected with PMSG (1,750 IU) and hCG (1,500 IU), 78 h apart, to induce oocyte maturation and ovulation. After hCG injection, the animals were artificially inseminated and sacrificed 30 to 36 h or 54 to 56 h later, and fertilized eggs were harvested from their oviducts. The recipients were synchronized and ovulation-induced by the same methods except that a 12-h delay was used, the dosage of PMSG and hCG was reduced to 1,500 IU and 1,250 IU, respectively, and insemination did not occur. When the fertilized eggs arrived at a nearby laboratory, CRISPR/Cas9 RNA was microinjected into the cytoplasm; the eggs were then surgically transferred to the oviduct of a recipient from the end of the infundibulum by exposure of the uterine horn and oviducts within 3 to 4 h. The recipients were raised normally but treated with special care, particularly during farrowing.

### Pig embryo manipulation and microinjection

The recovered newly fertilized eggs were centrifuged at 15,000xg for 10 to 15 min at 25°C to expose their pronuclei. The pronuclear embryos were added to a 20 μL microdroplet of D-PBS in a glass slide chamber and covered with mineral oil. The micro-manipulation was conducted under an inverted DIC (differential interference contrast) microscope at 200 to 300 x magnification. Each embryo was held in the proper position to reveal the pronucleus, and a mixture of single-guide RNA directed against two sites (sgRNA, 10 ng/μL each) and Cas9 RNA (70 ng/μL) was microinjected into the cytoplasm near the pronucleus using a capillary needle with steady flow.

### Construction of CMAH gene-specific sgRNA knockout and Cas9 vectors

In most genes, the 5’-end sequences usually encode the most important or the largest domains. The codon region of the porcine CAMH gene includes 14 exons; exon 1 contains 8 bp, and exon 2, which is 204 bp in length, is the largest exon (Fig 1A, large capital letters shaded in yellow). After verifying the sequences of exon 2 and introns 1 and 2 of the CMAH gene, we chose two GN_19_NGG Cas9 specific sequences; one of these has a sense strand site on exon 2, and the other has an antisense strand site located on intron 2 (Fig 1A, characters underlined in red). According to the sequences of the selected sites, two synthetic DNA primer pairs (shown in Table 8) were annealed as double-stranded DNA fragments, digested with BsalI and cloned into the ppU6-(BsaI)_2_-sgRNA vector [48]; in this way, two sgRNAs, ppU6-(CMAH ex2)-sgRNA and ppU6-(CMAH in2)-sgRNA, were constructed. Cas9 in the pCX-Flag_2_-NLS1-Cas9-NL-S2 vector was constructed by Su et al. [48]. To make it possible to use the RNAs for gene editing, the U6 transcription promoter of both CMAH ppU6-(CMAH ex2)-sgRNA and ppU6-(CMAH in2)-sgRNA was replaced by SP6, and CX in pCX-Flag_2_-NLS1-Cas9-NL-S2 was exchanged for the T7 promoter. To prepare sgRNA and Cas9 RNA for use in microinjection of porcine fertilized eggs, the RNAs were subjected to in vitro transcription using the MEGAscript^®^ T7 Transcription Kit (AM1334, Carlsbad, CA, USA) and purified using the MEGAclear™ Transcription Clean-Up Kit (AM1908, Carlsbad, CA, USA).

**Table 8.**
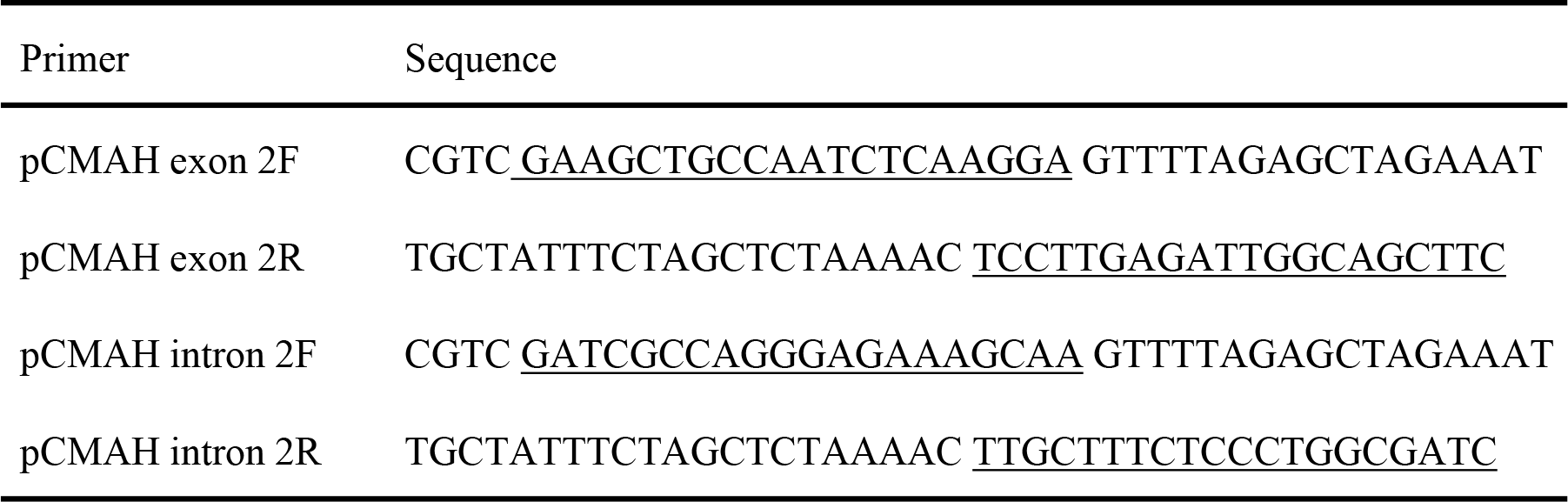
Primer Pairs Used To Construct sgRNA Expression Vectors.

### Screening of CMAH gene mutant pigs

Genomic DNA of all pigs delivered from foster dams or founders was isolated from tissue obtained from the piglet’s tails and purified using a genomic DNA purification kit (Fermentas/Thermo). The CMAH mutant pigs were first screened by PCR using 0.1 μg of genomic DNA and 0.25 ◻M each of CMAH Ex2 F (TGG AGC TGT CAA TGC TCA GG) and CMAH Ex2 R (TCA GAG AGC TGC CGT AAA GG) primers annealed at 55°C. Wild-type or site-mutated pigs produced an ~439-bp amplicon, and biallelic simultaneously mutated animals displaying the 161-bp deletion produced an ~278-bp amplicon. For further confirmation, all PCR products were verified by PCR product-direct sequencing (PDS) and PCR product/TA cloning/sequencing (PTS); from the latter, at least 6 colonies were picked and sequenced. DNA primer synthesis and DNA sequencing were conducted by Mission Biotech Ltd. (Taipei, Taiwan). The sequencing data were analysed using BioEdit software.

### Analysis of NGNA/NANA by HPLC

Samples of ear, tail and small intestine weighing approximately 100 mg were cut into small pieces in MQ water and incubated at 95°C for 30 min. After the samples had cooled to room temperature, 0.5 M H_2_SO_4_ was added to a final concentration of 25 mM. The mixtures were incubated at 80°C for 1 h to release the sialic acids from the samples. After centrifugation, the supernatant was collected, an equal volume of DMB (1,2-diamino-4,5-methylenedioxybenzene, Sigma-Aldrich, Inc.) solution (1.6 mg DMB in 1 mL of 1.4 M acetic acid, 0.75 M 2-mecaptoethanol and 18 mM sodium hydrosulfite solution) was added, and the mixture was incubated at 80°C for 2 h to label the sialic acids. The labelled NGNA and NANA used as standards were prepared as 1 mg/mL solutions and reacted under the same labelling conditions. The DMB-labelled sample was injected onto a Waters™ HPLC system (Waters 2475 Multi-wavelength Fluorescence Detector, Waters 717 plus Autosampler and Waters 600 Controller) with the Discovery^®^ BIO wide Pore C18 (5 μm, 4.6 × 25 cm) column. The analysis was performed using an isocratic mobile phase of methanol:acetonitrile:H_2_O (7:9:84) at a flow rate of 0.6 mL/min; the fluorescence detector was set at an excitation wavelength of 373 nm and an emission wavelength of 448 nm.

### PEDV challenge

#### Piglet treatment and facility

All CMAH KO neonatal piglets were delivered from three F0 female founders that were served by the male F0 founder; thus, all founders were half or full sibs. All founders were biallelic CMAH mutants carrying a biallelic 161-bp deletion (D/D type) or one allele deleted and the other mutated (D/M type) genetic background. The D/D type and/or D/M type piglets were used as described in the experimental section. The control piglets were non-gene-edited piglets that were concurrently delivered from wild-type sows at the same farm.

PEDV challenge was conducted in a negatively air-conditioned animal facility at the NPUST. The pens were equipped with stainless mesh floors that allowed the faeces to drop down to a collection plate. The room temperature was set at 30°C, and each pen was equipped with two extra electric power bubs.

During 4 h shipping, the piglets were kept at 25°C in dark containers. When they arrived at the challenge room, the mutant and wild-type piglets were grouped and placed in different pens. Approximately one hour later, all piglets were oral inoculated with PEDV, which diluted in commercial baby milk powder that had been reconstituted with warm drinking water. In experiments I and II, the animals in each pen had free access to 200 mL of fresh prepared baby milk and clean tap water that was changed every 4 h. In exp. III, PEDV was diluted with KO or wild-type sow’s milk obtained 2 days after parturition, and no milk was offered; instead, fresh drinking water was offered and changed every 4 hours. Other treatments were as described in experimental design III.

### Preparation of PEDV virus for Use in PEDV Challenge

New variant-PEDV (nv-PEDV) was isolated from a field case that occurred at Jimei farm in Yunlin County in central Taiwan in February 2015. Almost all of the affected one-week-old piglets died of watery diarrhoea. The aetiology of the disease was confirmed to be a virulent strain of PEDV (it was thereafter designated the Jimei strain); the sequence of this strain is almost identical to that of the strain that caused the epidemic outbreak of PEDV in the US in 2014 [36]. Although nv-PEDV can replicate in the Vero cell line, the nv-PEDV used in the challenge was prepared by oral inoculation of new born piglets that had not received colostrum to maintain its pathogenicity. The piglets were raised in a warm isolated chamber and were hand-fed fresh milk every six hours. Diarrhoea began to occur at 16-20 h after viral inoculation. The piglets were sacrificed 16-24 h after the observation of diarrhoea symptoms. The small intestinal content was collected by injection of 50 mL of DMEM supplemented with 10x P/S into the lumen followed by massage and extrusion from one end to the other end. The intestinal content was filtered through stainless mesh to clarify the content. Finally, the sample was centrifuged at 3000xg to precipitate all cellular debris, and the supernatant was collected and divided into 5-mL portions in sterile conical tubes. Three small fragments of intestine were subjected to paraffin-embedded tissue sectioning and IHC to confirm the presence of PEDV in intestinal epithelial cells (TGEV and rotavirus detection was also performed, and both tests were negative). A TCID_50_ was used according to standard virological methods to determine the viral content of the Jimei PEDV virus preparation used in the challenge study. The virus was maintained at −80°C until the challenge study was performed. Inoculation of the animals with PEDV was conducted as described by Jung et al. [7]. In brief, 10^3^ TCID_50_/mL of frozen nv-PEDV stock was thawed at hand temperature, and 9 mL of the thawed stock was mixed with 90 mL of reconstituted commercial baby milk or sow’s milk by repeatedly inverting the container. The CMAH mutant and wild-type piglets were inoculated with 10^3^ TCID_50_/10 mL PEDV orally by hand using a syringe.

### Experimental design

#### Exp. I: Challenge of 2-day-old neonatal piglets with nv-PEDV

In total, 6 D/D type and 6 wild-type piglets were used for PEDV challenge, and one D/D type and one wild-type piglet without virus treatment were used as controls; the latter were not housed with the infected piglets. All neonatal piglets were nursed for approximately 20 h to allow intake of colostrum and then delivered to a negatively air-conditioned facility.

#### Exp. II: Challenge of 3-day-old neonatal piglets with nv-PEDV

In this trial, 8 D/D and 1 D/M type mutant piglets and 9 wild-type piglets were used for PEDV challenge, and one D/D mutant piglet and one wild-type piglet without virus treatment served as controls. All of the neonatal piglets were nursed for approximately 44 h to permit intake of colostrum and dam’s milk. The detailed conditions of the PEDV challenge were the same as those used in experiment I.

#### Exp. III: Challenge of 2-day-old neonatal piglets with nv-PEDV followed by extended feeding of sows’ colostrum

In this trial, 9 D/D and 3 D/M type mutant piglets and 12 wild-type piglets were used for PEDV challenge, and one D/D type piglet and one wild-type piglet without virus treatment served as controls. All neonatal piglets were nursed for approximately 20 h to permit intake of colostrum and then delivered to a negatively air-conditioned facility. In this trial, the piglets were orally inoculated with PEDV as in experiment I and II and were not fed commercial baby cow milk; instead, they were fed their dams’ or other founder’s milk that had been collected within 20 h. From 4 to 24 h post PEDV inoculation (hpi), the piglets were fed 20 mL of sow’s milk by hand every 4 h; whole milk was fed at 4 and 8 hpi, and skim milk was fed from 12 to 24 hpi. From 24 hpi to 72 hpi, 20 mL of lactated Ringer’s solution supplemented with 5% glucose was fed to each piglet every 4 h. The piglets were randomly allocated to sacrifice at 24 hpi (3 piglets), 48 hpi (3 piglets), or 72 hpi (6 piglets), and the small intestines were sampled.

### Clinical observations

After inoculation with PEDV, the piglets’ behaviour, including vomiting, diarrhoea, and lethargy, was observed and recorded every 4 hours for 3 days. When the piglets died or at the end of the experiment, their body weights were recorded, and they were necropsied on the same day.

### Sampling

The intestines of all piglets were sampled at the upper and middle region of the jejunum and the upper part of the ileum by resecting a portion of the intestine approximately 10 cm in length. This piece was then ligated at both ends with surgical string, cut down, and a suitable amount of 10% formalin was injected into the luminal space. The entire sample was then immersed in ~15 mL 10% formalin and fixed for at least for 24 hours.

### H/E and immunofluorescence (IF) staining

After fixation, the samples of intestine obtained from the piglets were sliced, embedded in paraffin, and sectioned at 3 to 4 μm thickness. The sections were placed on slides, de-waxed in xylene and sequentially treated with 100%, 95%, 80% and 70% ethanol; the slides were then stained by H/E. For IF staining, the slides were de-waxed in xylene and 100% ethanol and further heated in boiling TAE buffer for 3 min to activate the antigen. After cooling to room temperature, the slides were washed with PBS for 15 min, and the tissues were stained with a primary antibody against PEDV (prepared by Dr. CM Chen) and a commercial secondary antibody, FITC-conjugated goat anti-mouse immunoglobulin (Cappel). After immersion in DAPI solution, the slides were sealed with 10% glycerol, and the signals were observed on an Olympus BX50 microscope (Olympus, Japan) enlighten by UV-light.

### Pathology evaluation

The criteria used to score immunofluorescence (IF) staining and histopathological lesions (I/H score) associated with PEDV are shown in Fig 8. PEDV mainly infects the epithelial cells that form the mucosa of the small intestine. In the early stage of PEDV infection, only IF staining allows us to observe whether or not epithelial cells have been infected by PEDV. Therefore, at that stage, the percentage of IF-positive cells was the only criterion used to determine the severity of PEDV infection. However, in the middle to late stages of infection, the severity of PEDV infection is better judged by the degree of villar atrophy because infected cells often defoliate from the mucosa and IF may not reveal the PEDV-infected cells. Therefore, the lesions were scored from 1 to 5 as shown in Fig 8C; the scores combined the results of both IF staining and histopathological inspection in an I/H score that was used in the final statistical analysis.

### Statistical analysis

All of the clinical and viability data were recorded and analysed using GraphPad Prism 6 (GraphPad Software, Inc.). The survival rate (curves) of the piglets after PEDV challenge was analysed using the Log-rank (Mantel-Cox) and Gehan-Breslow-Wilcoxon tests. The t-test was used to analyse the body weight and immune/histopathologic data obtained from the intestinal samples from all experimental piglets. The significance level (*) was set at 0.05.

## Acknowledgements

The authors would like to express their sincere thanks to Mr. Chi-Yun Hsu, Mr. Shau-Ching Hseu and Mr. Ci-Hong Wong for assistance with the care of the experimental animals, particularly that of the KO pigs. Thanks are also expressed to Ms. Ming-Shing Liu for technical assistance with pig surgery and embryo transfer and to Dr. Chao-Nan Lin for his assistance with the PEDV challenge trial at the National University of Science and Technology.

## Supplementary Information

**S1 Fig. Analysis of CMAH Gene-Edited Offspring from the First Parity.** A. The PCR products that revealed more than one band were further subcloned into the TA vector for colony purification and sequencing. B. Offspring with mutations at site I (exon II) and site II (intron 2). C. The offspring carrying two sites mutated simultaneously, with deletion of a 161-bp DNA fragment, and some of them showed further indel of +1 or −5 bp.

**S2 Fig. Analysis of CMAH Gene-Edited Offspring from the Second Parity.** A. The PCR
products that revealed more than one band were further subcloned into the TA vector for colony purification and sequencing. B. Offspring with mutations at site I (exon II) and site II (intron 2). C. The offspring carrying two sites mutated simultaneously, with deletion of a 161-bp DNA fragment, and some of them showed further indel of +1 or −5 bp.

**S3 Fig. Analysis of CMAH Gene-Edited Offspring from the Third Parity.** A. The PCR products that revealed more than one band were further subcloned into the TA vector for colony purification and sequencing. B. Offspring with mutations at site I (exon II) and site II (intron 2). C. The offspring carrying two sites mutated simultaneously, with deletion of a 161-bp DNA fragment, and some of them showed further indel of +1 or −5 bp.

**S4 Fig. HPLC Analysis of NGNA/NANA in the Ear Tissues of Six F1 Offspring of the CMAH KO Founders. T**he retention times of NGNA and NANA are shown as numbers on the peaks.

**S5 Fig. Gross Appearance of the Small Intestine of Neonatal Piglets at 72 hpi or at the Time of Death during the Trial.**

